# Protein condensate fusion mapping using Hippopede curve

**DOI:** 10.1101/2023.10.12.562122

**Authors:** Sumon Sahu, Jennifer L. Ross

## Abstract

Protein and nucleic acid condensation in cells has been a major focus of biophysical studies in recent years. Such droplets, condensates, or membraneless cellular organelles that form by condensation have been shown to behave like liquid or viscoelastic fluids. In vitro reconstitution of condensates of purified protein and nucleic acid have been used to better understand the phenomena, specifically to quantify the material properties of these condensates. One of the signature properties of a liquid-like condensate is the fusion of two round droplets into a larger droplet. Quantitative imaging and analysis of the fusion process can report the biophysical properties of the condensates, such as surface tension and viscosity in a non-invasive way. Here we take advantage of a model in soft matter physics, hippopede curves, to the condensate fusion problem to estimate these biophysical properties directly and compare the results with existing models in the field. In addition, this method approximates additional parameters such as the surface area and surface curvatures during the fusion process over time.

## Introduction

Phase separation has emerged as an universal method for assembling biomolecules into membraneless compartments that perform crucial biological functions inside cells such as transcription, genome organization, stress and immune response, neuronal synaptic signalling, intracellular organization, and others(1–4). Dysfunctional condensates formed by phase separation have been linked to pathological aggregates in diseases such as cancer, neurodegenerative diseases, and viral infections (2, 4, 5).

The interface of a condensate or membraneless organelles is what determines the “inside” from the “outside” of the droplet. The interfacial tension of phase separated droplets is critical to maintain the shape, concentration, and molecular exchange across the interface in the condensate. The surface tension of condensates is also capable of doing work to move and organize intracellular systems through capillary forces and wetting (6). These interactions have been proposed to drive biological processes, such as RNA splicing at the surface of nuclear speckles (7). Thus, the surface tension is an important, biologically relevant material property of membraneless organelles to measure.

Recapitulation of protein and nucleic acid condensation in vitro has allowed direct probes of the material properties of these condensates through a variety of techniques. In vitro, it is easy to access these condensed droplets to directly probe the surface tension. For example, biophysical techniques have been employed such as micro-pipette aspiration (8), microrheology and active fusion through optical tweezers (9–12), shear relaxation (13), sessile droplet shape analysis (14), and passive coalescence (15, 16).

Measuring the surface tension of protein condensates in live cells poses various challenges. The cytoplasm and nucleoplasm are viscoelastic environments full of polymers that can cage the droplets. These environments are also active, being rearranged by the enzymes acting on the polymers. This non-equilibrium, crowded environment can change the condensate and the condensate and alter the environment (17, 18) The well-used method for measuring the surface tension of droplets in live cells is to analyse passive coalescence and measure the aspect ratio of a bounding ellipse over time (15). This method is restricted to measuring the fusion of same-sized droplets and provides little information about the surface dynamics.

Given the importance of understanding the material properties of condensed droplets, and the need to extract more measurable information about the merging process, we have developed a quantitative image analysis method to deduce the surface properties of droplets including surface area, curvature, and surface tension. Here, we use spinning disc confocal imaging to create 3D reconstructions of droplets during merging events in vitro. The boundary of the droplets is determined and modeled using the hippopede curve solution to model the shape, calculate the surface area, and determine the curvature of the protein condensates as they fuse, frame by frame. This method was adopted from analyses used for soft matter systems, such as coascervates, aqueous two-phase systems, and emulsions (19, 20).Finally we calculate the viscosity and surface tension and compare our results with the ellipse fit method that is used in the condensate field.

## Experimental Methods

### Reconstituted condensation in vitro

The protein purification and imaging chamber design were the same here as previously reported (3). Briefly, MAP65 and MPA65-GFP plasmids were gifts from am Dixit (Washington University, St. Louis) and purified as described (21, 22). MAP65 is expressed in *Escherichia coli* BL21(DE3) cells, grown to an OD_600_ of 0.6-1. Cells are lysed with tip sonication, and the lysate is clarified by centrifugation. The supernatant is incubated with Ni-beads with affinity for the 6*×* histidine tag and eluted with imidizole. Purified MAP65 was buffer exchanged to PEM80 (80 mM PIPES, 1 mM EGTA, 2 mM MgCl_2_) by dialysis, aliquoted, and stored at −80 ^*o*^C. For purification and concentration measurements the purified MAP65 was checked on an SDS-PAGE gel and Nanodrop. MAP65 with 10% MAP65-GFP were combined in PEM80 and incubated at room temperature for 20-25 min. A 10 *μ*l flow chamber made using glass slide (Fisher), silanized coverslip, and double sided tape (3M) was coated with 5% F127 in water and incubated for 7-10 min. The sample was flowed into the chamber, and the chamber was sealed with epoxy on both ends. The chamber was kept cover slip-side down for 15 min in order to let the condensates settle by gravity. The coalescence videos are taken immediately after the 15 min incubation.

### Confocal imaging

Imaging of GFP-MAP65 condensates is performed using spinning disc microscopy (Yokogawa CSU-W1 50um Pinhole) on an inverted Nikon Ti-E microscope with Perfect Focus and 100x oil immersion objective (1.49 NA) imaged onto a Andor Zyla CMOS camera. Image acquisition is performed with 488 laser. The image scale is 65 nm per pixel. Images are displayed and recorded using the Nikon Elements software. Time series data was saved as .nd2 files with metadata and analyzed using ImageJ/FIJI and MATLAB.

### Pre-processing and image analysis methods

The images are pre-processed by image analysis routines to reduce noise and extract relevant information. First, a median filter of 4*×*4 matrix is applied on the raw images in figure 2(a) to smooth the images, which reduces noises. Then the images are binarized in MATLAB. After that, an appropriate area size filter, ‘bwareafilt,’ was applied to eliminate the areas that were not of interest. The centroids of the condensates in the images are translated to the image centers and rotated according to their orientation, in order to keep the coalescence process on a horizontal axis, facilitating the fitting routines. We use MATLAB’s built-in function ‘regionprops’ to quantify the area, centroid, the orientation of the droplets in the images. Finally, the boundary/interface of the condensates is detected by the canny edge detection algorithm in MATLAB (Fig. 2(b)).

To convert the binary boundary map into a polar (*r, θ*) plot, we introduced a square mask around the centroid to store *R* and Θ variables, using MATLAB’s mesh grid and ‘car2pol’ functions. Initially, the centroid of the dumbbell shaped structure in the coalescence process serves as the origin for these new matrices. However, as the schematics depict in Fig. 1(ii), the origin of the model is not always the centroid of the evolving shape. Therefore, we sampled the origin slightly left to right to find the origin location that fits the model best. Each time, according to the model parameters, a cost curve was generated to serve as a metric to evaluate the goodness of fit for that location. We picked the minima of the cost curve, which reports the origin according to the model and provides as the best fit or agreement to the model.

**Figure 1.**
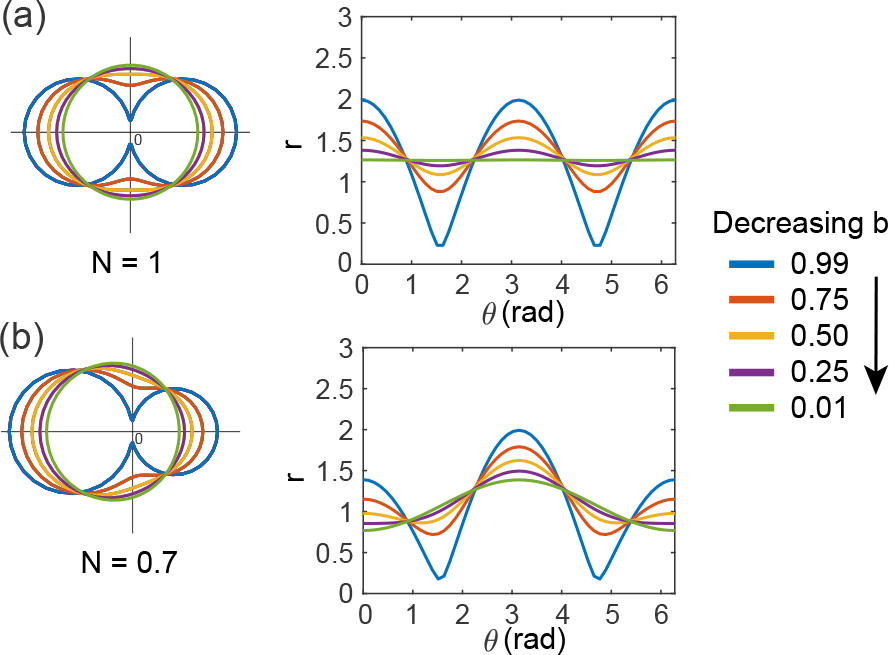
Contours calculated from *r*(*θ*) from eqn. 1 is shown for two scenarios, (a) circles of same radius or same size with N=1, (b) circles of different radius or heterogeneous in size with larger left circle, N=0.7. Here the color represents different coalescence parameter b values, from blue to green indicating fusion progression.

## Results

### Shape analysis of condensate coalescence in vitro

The sample chambers were coated with a triblock copolymer (Pluronic F127) so that the droplets do not wet the surface with a high contact angle, as previously described (3). Given the axial symmetry of the fusion process, the 3D droplets that sit on the chamber bottom surface can be projected down to a plane around the axis. The captured confocal cross-sectional image through the mid-plane of the condensate represents the 2D plane projection (Fig. 1(a)). The surface boundaries of the droplets are extracted for all frames by binarizing the image and applying the canny edge detection algorithm in MATLAB (see Experimental Methods).

When two liquid-like protein condensates fuse into one condensate, there are two cases two consider: (1) the condensates are the same size or (2) the condensates are heterogeneous in size (Fig. 1(b)). The generalized hippopede curve approach, an empirical geometrical model, can capture both of these scenarios to model the shape evolution during the process in 2D polar coordinates, (*r, θ*). Prior work has used the hippopede curve to approximate the surfaces of two equal-sized coalescing spheres (23). A modified version of that model has been used to incorporate unequal size spheres (19, 24). Written in polar coordinated, the contour function is:

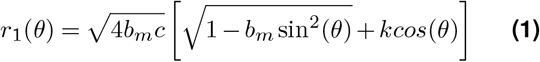

where 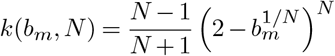

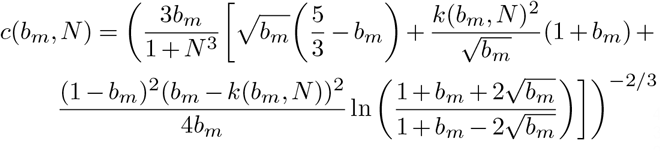

In these equations, *b*_*m*_ and *N* are independent parameters. The parameter 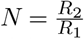 is the ratio of radii of two spheres before the initial point of contact, where *R*_1_ is the radius of the left sphere and *R*_2_ is the radius of the right sphere. The coalescence parameter, *b*_*m*_, carries a value from 1 to *b*_*min*_ that describes the degree of coalescence (Fig. 1(b)), where *b*_*m*_ = 1 corresponds to the initial scenario when two individual spheres are just in contact, and *b*_*m*_ = *b*_*min*_ corresponds to the final state of complete coalescence to a complete sphere. The *b*_*min*_ depends on *N*. The value of *N >* 1 and *N <* 1 change depending on the size inequality and position of the spheres (Fig. 1(b)). For this work, we always position the larger sphere on the left, so *N* ≤ 1 for our purpose, but this is not required for the method to work. The second *k*-term in Eq. (1) is a function of *b*_*m*_ and *N* and is modified to *c*(*b*_*m*_, *N*) to allow for two unequal size radii in the problem. From prior work, the shapes from this model were validated through finite element methods (25) and boundary element methods (26).

The geometrical model in Eq. (1) is a scale-free equation, which assumes the radius of the left sphere is 1 as a reference. We restricted ourselves to *N* ≤ 1 and introduced a scale factor, *S*, as a parameter that scales the dimensions of the left sphere accordingly while fitting the experimental data. Hence, the final equation we use is *r*_*m*_(*θ*) = *Sr*_1_(*θ*) to re-scale and fit the experimental data *r*_*e*_(*θ*) in each frame. Each frame from a condensate fusion time-series is rotated accordingly so that the fusion occurs along the x-axis (Fig. 2(a),(b)). The surface contour positions from the binary images are extracted as *r*_*e*_(*θ*) values to fit the model to get the parameters *b*_*m*_, *N*, and *S* (Fig. 2(c). Since, *N* and *S* should be constant and dependent on each other for a given pair of droplets, we calculate *S* from radius of larger droplet, *r*_1_, on frames before fusion and use that *S* to fit *r*_*e*_(*θ*) using *b*_*m*_ and *N* .

In the hippopede curve model, while the fusion process is ongoing, the condensate neck is considered to be the origin. Since the merging condensate shape changes during the fusion process, the location of the origin with respect to the merging droplet boundaries shifts over time. We observed that finding the (*r, θ*) coordinate origin has an impact on shape fitting. Therefore, we sample a few pixels to the left and right from the centroid of the merging droplet to find the origin that fits best to the *r*_*e*_(*θ*). The best fit is calculated from minimization of a cost function defined as 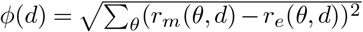

The hippopede curve model fits the raw image contours well after finding the origin with minimum cost using the [*b*_*m*_, *N, S*] parameter optimization (Fig. 2(d-f)). For every *b*_*m*_ and *N* we the calculate cost function and finally find the minima and corresponding *b*_*m*_ and *N* (Fig. 2(f)).

**Figure 2.**
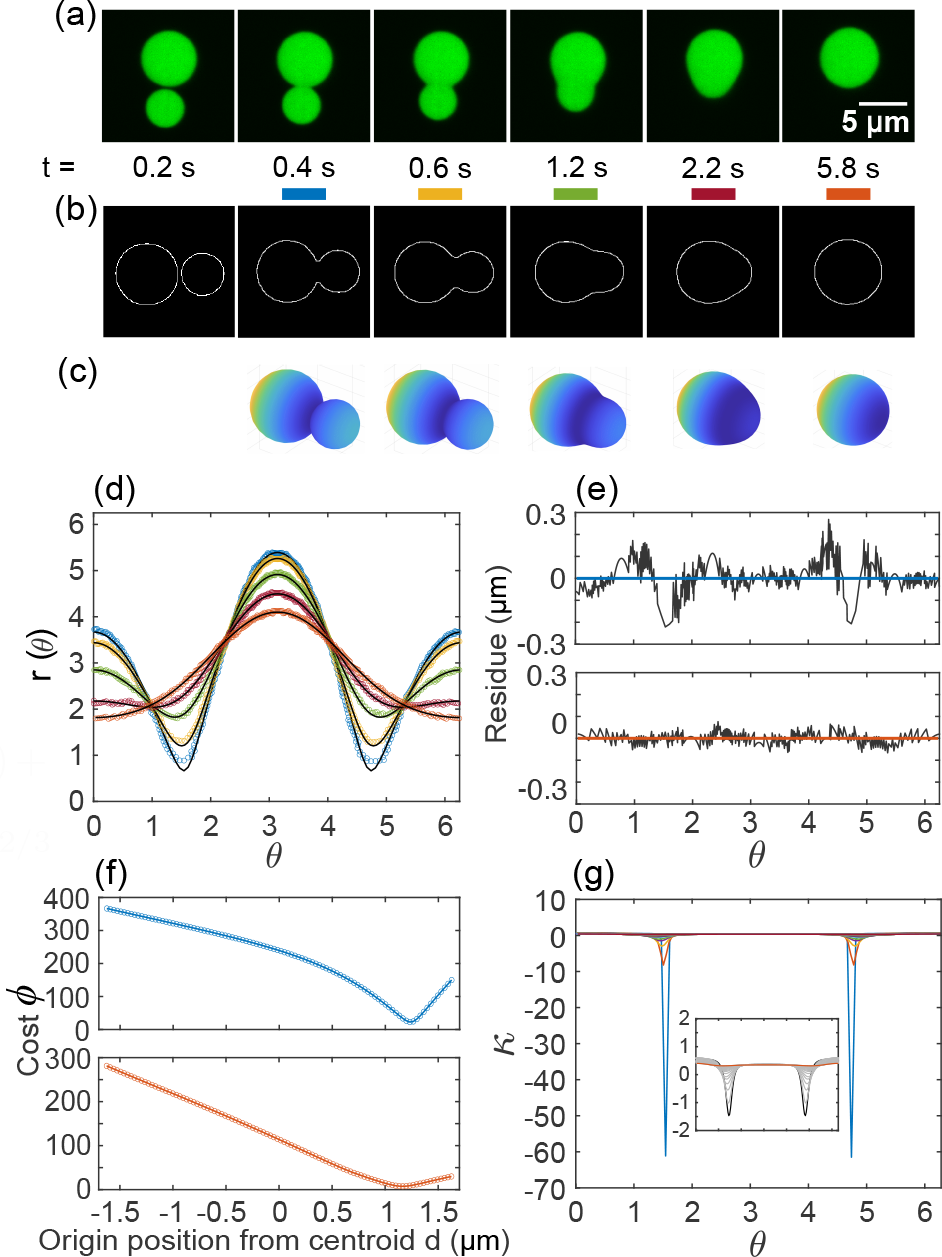
Geometrical model of the condensate interface evolution: (a) Time series of a coalescence event of two condensates is shown over time with 200 ms between frames and *t* = 0 s indicates the first frame of the time series (not shown). Scale bar is 5 *μm*. (b) Interface maps of the corresponding images are shown in white with a black background. Raw images are first translated and then rotated to the horizontal axis according to the orientation of the condensates. (c) 3D surface of revolution as predicted from the hippopede model fit to the data from the corresponding frames. (d) The *r − θ* plot for individual frames: 0.4 sec (blue), 0.6 sec (yellow), 1.2 sec (green), 2.2 sec (red), and 5.8 sec (orange) are shown with data from interface map. The corresponding model fits using the scaled version of the equation Eq. (1) are shown in black. (e) The residuals of the data and the fit are plotted for *t* = 0.4*s* (blue) and *t* = 5.8*s* (orange) to show goodness of fit. The black lines are fits compared to colored experimental data. (f) The cost functions 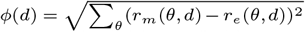 are plotted against origin distance from centroid, *d* (origin sampling) for *t* = 0.4*s* (blue) and *t* = 5.8*s* (orange). (g) The curvature, *κ*, of the interfaces calculated from the model fit parameters as the fusion progresses. Time frames: 0.4 sec (blue), sec (yellow), 1.2 sec (green), 2.2 sec (red), and 5.8 sec (orange). Inset shows calculated curvatures from model for subset of frames *t* = 1*s* (black) to 5.8*s* (orange) with 0.2*s* apart in-between frames (grey).

### Using shapes to estimate surface mechanics

In a complete coalescence process of condensates, surface tension coupled with curvature gradients drive the fusion process. At the initial point of contact, the process starts with a divergence in curvature at origin (Fig. 2). This sharp feature smooths out as the fusion progresses (Fig. 2). To calculate curvature smoothing over time from the model fits, we estimate curvatures at each time point using the model fit from the experimental images. In polar coordinates (*r, θ*), curvature *κ* takes the form,

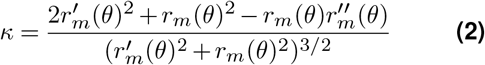

In this equation, the primes indicate derivatives with respect to *θ*. Using Eq. (1) with fit parameters, we calculated the curvature at each time point. Indeed, the calculated curvatures show the reduction of sharp features over time (Fig. 2(g)). As a result, the model finds the curvature of the final droplet to be *κ* = 0.34 *±* 0.02 1*/μm*. To double-check if the reported curvature from the model was correct, we measured the radius of the final condensate in ImageJ. We found *κ ∼* 0.339 *±* 0.002 1*/μm* (n=5) using the equation 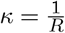, which is a good comparison.

Using our hippopede fits, we can measure the dynamics of the coalescence event over time. The parameter, *b*_*m*_, serves as a measure of the shape for the coalescence, which we call the coalescence parameter. We extract this parameter from best fits to the experimental image over time and find that the *b*_*m*_ relaxes exponentially to *b*_*min*_ with a characteristic time, *τ*_*b*_ = 2.10 *±* 0.19 s. The image frame at *t* = 0.4 s indicates the initiation of the fusion process, the frames before that time have images where the model is unsuitable for fitting. Thereby, these frames were excluded from the exponential fit (Fig. 3(a)).

**Figure 3.**
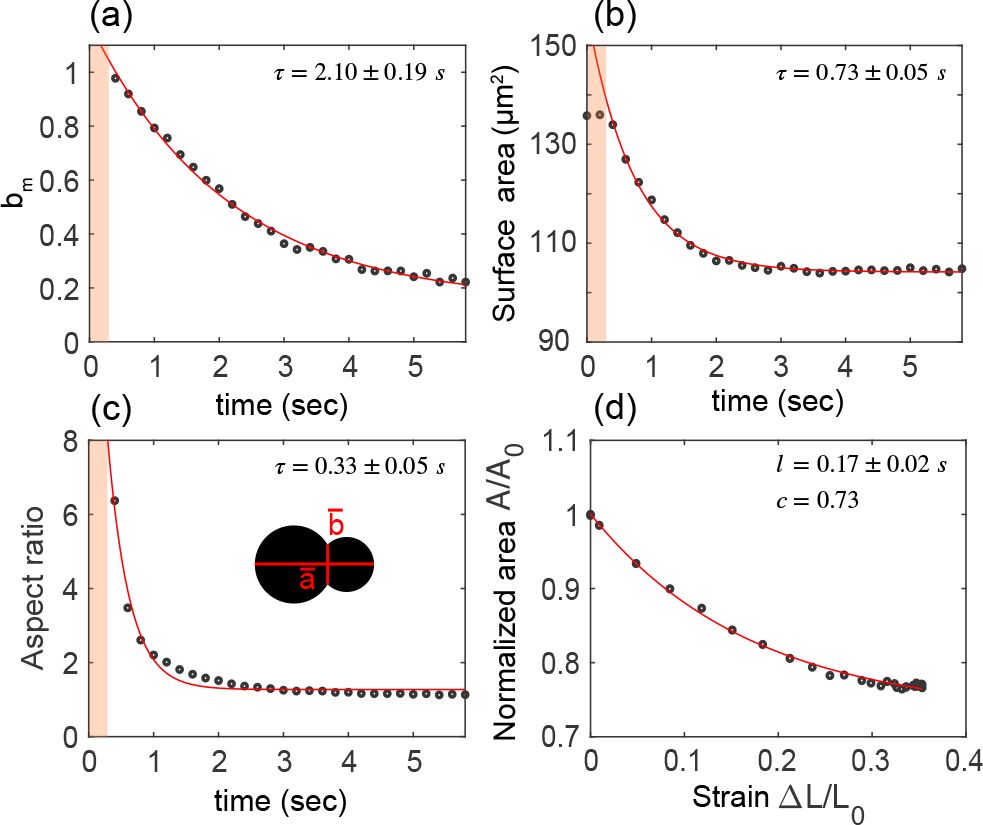
Dynamics metrics for coalescence events over time. (a) The coa-lescence parameter, *b*_*m*_, in the model is plotted with time for the coalescence process and fit with an exponential decay curve with *τ*_*b*_ = 2.10 *±* 0.19 *s*. (b) The surface area change during coalescence is calculated from the model fit parameters and shows an exponential relaxation. The exponential fit yields *τ*_*s*_ = 0.73 *±* 0.05 *s*.(h) The aspect ratio, 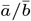, calculated from the model is plotted with time. The exponential fit yields *τ*_*model*_ = 0.33 *±* 0.05 *s*. The shaded red zones in (a), (b), and (c) indicate the time points where condensates are separated. (d) The normalized area is plotted as a function of strain during droplet coalescence and fit with an exponential. The characteristic strain turns out to be *l ∼* 0.17 with final normalized area from exponential fit to be 0.73.

Since the shape of the condensate has a rotational symmetry around the x-axis, we can construct a surface of revolution from the model with best-fit parameters to create a three dimensional model of the condensates while merging (Fig. 2(c)). From this, we can calculate the change in surface area during the merging process. The surface area of two droplets before fusion initiation were estimated from the binary images. First, the radii of the condensates were approximated from the areas calculated by MATLAB regionprops function. Then the surface area *A* = 4*πr*^2^ was calculated from those radii. For frames after the fusion initiation, the surface areas were calculated from the model parameters using Eq. (1) with *r*_*m*_(*θ*) = *Sr*_1_(*θ*). The boundary integral was performed numerically in MATLAB using

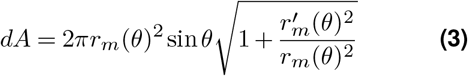

The *dA* is integrated over *θ* from 0 to *π* to calculate *A*. The surface area relaxes in exponential fashion with *τ*_*s*_ = 0.73 *±* 0.05 *s* (Fig. 3(b)). The surface area of the final frame of the condensate is calculated from model to be 104.8 *μm*^2^ (n=5). Using the final image, we can directly estimate the surface area from the measurement of the radius using ImageJ and find 108.7 *±* 1.4 *μm*^2^ which is near the model prediction.

Another parameter we can use to determine the dynamics is the ratio of the neck width 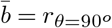 to the end-to-end distance 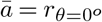, which represents aspect ratio for the hippopede curves: 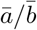 (diagram inset, Fig. 3(c)). These values are determined using the model fit for each image of the merging event over time. In the frames before the fusion, since 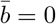, the aspect ratio values are not calculated as it diverged to an extreme value. The aspect ratio, 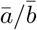, also shows an exponential decay with *τ*_*model*_ = 0.33 *±* 0.05 s (Fig. 3(c)).

When the droplets coalesce, the volume of two condensates remain constant (3), and the surface area decreases to a minimum value when there is a single sphere (Fig. 3(b)). During this process, an axial strain is developed. Depending on the physical state of the condensate, liquid, or viscoelastic, the final shape rearranges into spherical or a dumbbell shaped object. In case of liquid-like condensates, the normalized area traces out an exponential curve with respect to the strain for a given *N*. The final strain and normalized area for this particular fusion turns out to be 0.35 and 0.77 (Fig. 3(d)). If the fusion was arrested due to visco-elastic properties of the material the final shape would be found as a point on the curve.

It should be noted that when we perform the fitting, we can also fix the value of *S* from the [*b*_*m*_,N,S] fit to the initial frame when fusion just started instead of *r*_1_ estimated from frames before fusion, and allow the ratio *N* to vary. Moreover, we can also fix *N* from the beginning and allow *S* to vary. In order to determine if either method was better or if they were significantly different from one another, we repeated the fitting for the same data using these two approaches. Using the dynamics of the coalescence parameter as a metric, we find that there is no significant difference between the shape dynamics over time if we fix *S* and allow *N* to be fit (Fig. 4(a)), or if we fix *N* and allow *S* to be the fitting parameter along with *b*_*m*_ (Fig. 4(b)). We could not allow both N and S to be free parameters, as they are dependent on each other. The fits using single parameter *b*_*m*_ yielded poorer fits to the condensate contour.

**Figure 4.**
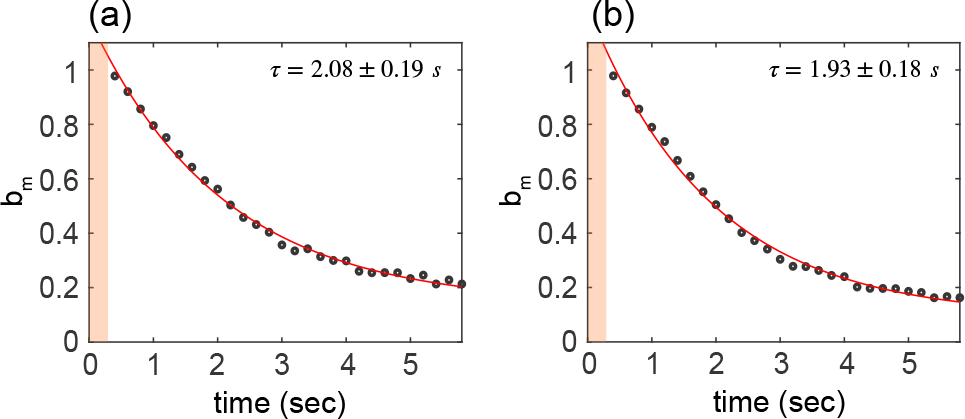
Comparison of two more approaches to using the hippopede model. The coalescence parameter, *b*_*m*_, over fusion process calculated from [*b*_*m*_, N] fit when *S* (calculated from first frame of fusion using [*b*_*m*_, N, S] fit) is kept constant in the model, fitted with an exponential decay curve yielding *τ*_*b*_ = 2.08 *±* 0.19 *s*. (b) The coalescence parameter, *b*_*m*_, over fusion process calculated from [*b*_*m*_, *S*] fit keeping *N* constant in the model is fitted with an exponential decay curve yielding *τ*_*b*_ = 1.93 *±* 0.18 *s*.

### Comparison of hippopede curve fitting and ellipse fitting

Using the hippopede curve approach, we use the droplet contours from the confocal images of droplet fusion to determine the shape dynamics during the fusion process, which gives us information on surface area and surface curvature. In addition, we can use this information to determine the surface tension of the droplets.

The most widely-used method to characterize the fusion process uses an ellipse circumscribed around the drops to each image as the droplets merge (Fig. 5(a),(b)). The aspect ratio, given by the semi-major axis length divided by the semi-minor axis length of the ellipse, is measured and plotted over time (Fig. 5(c). The aspect ratio relaxes over time and reports the characteristic time scale of the dynamics. When we use the ellipse method for the same fusion event (Fig. 2), the aspect ratio relaxation time we measure is *τ*_*ellipse*_ = 1.19 *±* 0.05 s (Fig. 5(c)). Comparing the characteristic time using the ellipse fitting method (Fig. 5(c)) to the characteristic times for the coalescence parameter, *b*_*m*_, the surface area, and the neck to length aspect ratio measured from the hippopede fits (Fig. 3), we find that the coalescence parameter has a characteristic time almost twice to that measured from the ellipse method. Interestingly, the relaxation of the surface area is actually faster than timescale estimated from the ellipse method for using the coalescence parameter.

**Figure 5.**
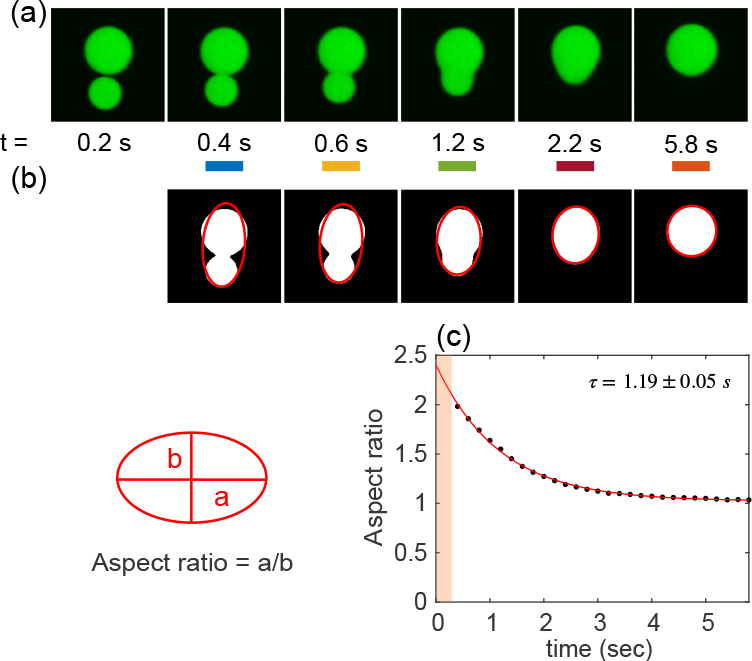
(a) Time series of the same coalescence event as before shown for t=0.2 s, 0.4 s, 0.6 s, 1.2 s, 2.2 s, 5.8 s. (b) Each image in the showed time frames are binary and fit to ann ellipse. From the fit major axis length a and minor axis length b are measured to calculate aspect ratio, a/b. (c) The aspect ratio relaxes as an exponential with time, with a characteristic timescale *τ* = 1.19 *±* 0.05 *s*

Using the same data sets of multiple coalescence events, we compare the hippopede curve fitting and circumscribed ellipse methods using data from both equal and unequal size condensate merging events. We compare characteristic length parameters from each method. Specifically, we compare the *S* parameter for the hippopede model with the characteristic length scale for the ellipse method, defined as: 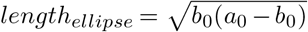 (15), where *a*_0_ and *b*_0_ are the semi-major and semi-minor axis lengths when the condensates are just in contact. We find that they are linearly related with a slope of 1.0 ± 0.2 (Fig. 6(a)). So, if we choose semi-major or semi-minor axis of the ellipse to calculate *length*_*ellipse*_ it should be nearly equal to *length*_*hippopede*_.

**Figure 6.**
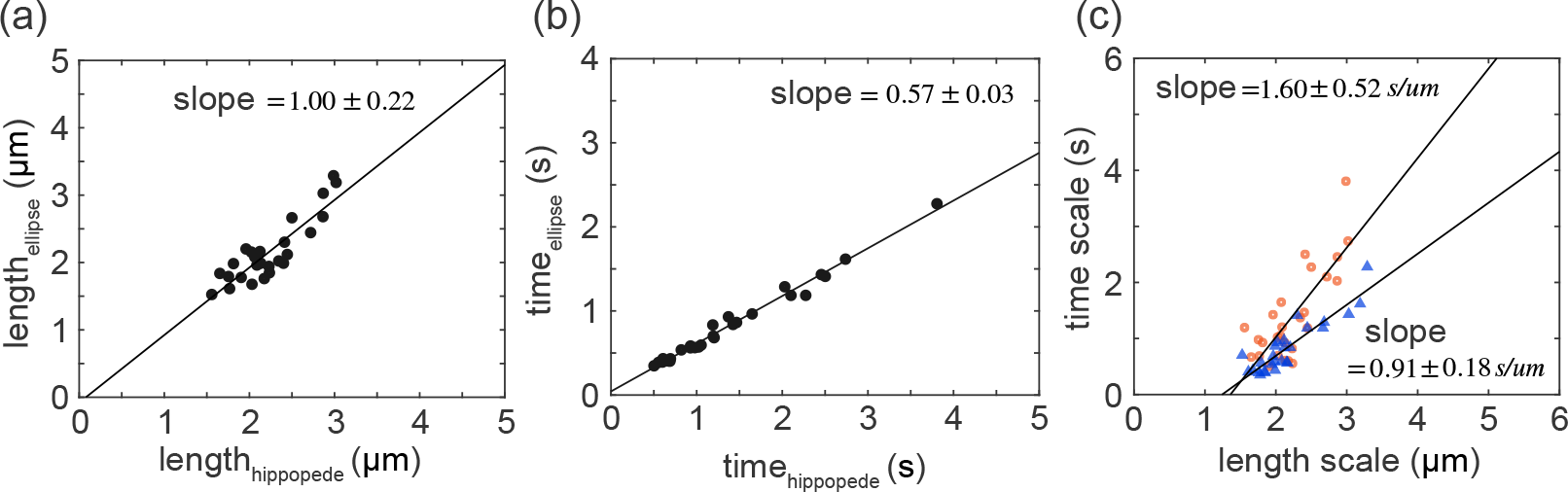
Length and time scale correlation between two methods: (a) The length scales *length*_*hippopede*_ and *length*_*ellipse*_ were calculated for same coalescence events by hippopede method and ellipse fit methods, respectively, and plotted for all data points (n=27). The data are fitted with linear trend with slope 1.00 *±* 0.22. (b) Similarly, all time scales calculated for same events are plotted and fitted with a linear fit that has a slope 0.57 *±* 0.03. (c) Finally, the calculated length scales and time scales calculated from two methods are plotted against each other. The hippopede fit approach (orange circles) has higher slope, *∼* 1.75 times higher hence larger inverse capillary velocity, compared to ellipse method (blue triangles).

Next, we compare the timescales between two methods *time*_*hippopede*_ (which is relaxation time of *b*_*m*_) with *time*_*ellipse*_ (relaxation time of ellipse aspect ratio) by plotting these time scales against each other (Fig. 6(c)). These time scales are linearly correlated with a slope 0.57 *±* 0.03 (Fig. 6(b)).

Taken together, we plotted timescale and length scale from these two methods, and calculate inverse capillary velocity. The inverse capillary velocity estimated by hippopede curve and ellipse fit method turns out to be 1.60 *±* 0.52 and 0.91 *±* 0.18, respectively (Fig. 6(c)). Thus, the estimate of the inverse capillary velocity is *∼* 1.75 *−* 2 times higher for the hippopede method compared to the ellipse fit method. Using the viscosity calculated from Stokes-Einstein equation (3), we estimate that the surface tension of liquid-like droplets is *γ*_*e*_ *∼* 5.2 *μN/m* from ellipse method and 1.75-fold lower, *γ*_*h*_ *∼* 2.9 *μN/m* for hippopede curve estimate.

Although the ellipse fit method is easy to use and provides quick output on droplet material state properties, its use is limited to merging events where the droplets are the same size, which limits the number of merging events that can be used. Moreover, the ellipse fit method does not quantify the time evolution of the surface area and curvature. Using the hipopede curve approach we were able to measure the changes in surface area and curvature evolution over time for unequal size condensates, as well as capture the length scales and timescales of the process. Also, using a single equation, we can create a 3D model reconstruction of the condensate fusion. We believe that the boundary shape modelling using the hippopede method is more rigorous and detailed analysis of condensate fusion.

## Discussion

We have shown that using a modified hippopede model, we are able to estimate the physical parameters of condensates during merging events to get estimates of the evolution of the surface area, surface curvature, and shear as well as the surface tension. This method can work for droplet merging with different sizes.

While the hippopede model fits the experimental data well and estimates mechanical and geometric parameter results, there are a few limitations. For instance, the hippopede model requires that the droplets start and end in spherical shapes. Unfortunately, some condensates inside a cell are not always round and may possess unique challenges as their morphology is unique and changes over time due to continuous interactions with neighboring structures. Other challenges of the fitting were overcome by using our cost function, such as identifying the neck of the merging event to position of origin of the curves, and estimating *b*_*m*_, *N*, and *S*.

Finally, we should mention that there are other shape-fitting methods that can be employed including using Cassini oval co-ordinates to estimate and model dumbbell shaped interfaces (27). This method has two fixed points or foci, accommodating for two droplet scenario before the fusion initiation step. In the generalized Cassini oval construction, exponents and expansions were used to incorporate unequal size nuclei (27, 28). Although we do not elaborate on it here, we have also tried to use Legendre polynomial expansion to fit unequal size MAP65 condensate during coalescence, but this method failed to fit high curvature frames and required more than four fit parameters to get a comparatively good fit with experimental data.

Future work can compare the surface tension estimates from hippopede model with measurements using micropipette aspiration or other mechanical probe techniques to determine if the model and direct measurement agree well.

## Author Contributions

SS performed the experiments, designed the analysis and wrote code to analyze the data. SS and JLR wrote and edited the article.

## Acknowledgments

We thank Chris Santangelo and Mahesh Gandikota for helpful discussions. This work was funded by a grant from the National Science Foundation NSF BIO-2134215 to JLR.

## Code Availability

Custom made MATLAB scripts created for this study will be available in a persistent repository upon publication.

